# Detection of rifampicin resistance in sputum samples by PCR-ELISA

**DOI:** 10.1101/2022.09.19.508626

**Authors:** Rosa Bellido-Pantoja, Edson Pacheco-Ascencio, Eddy Valencia-Torres, Omar Caceres-Rey

**Affiliations:** National Reference Laboratory of Biotechnology and Molecular Biology, National Center of Public Health, National Institute of Health. Chorrillos 15066, Lima, Peru; School of Biology, Faculty of Natural Sciences and Mathematics, Universidad Nacional Federico Villarreal. El Agustino 15007, Lima, Peru; National Reference Laboratory of Mycobacteria, National Center of Public Health, National Institute of Health. Chorrillos 15066, Lima, Peru; School of Human Medicine, Faculty of Health Sciences, Universidad Científica del Sur. Villa El Salvador 15067, Lima, Peru

**Author notes:** Corresponding Author, (O. Caceres). E-mail addresses (R. Bellido), (E. Pacheco), (E. Valencia).

**Keywords:** Tuberculosis, *Mycobacterium tuberculosis*, Rifampicin, Resistance, PCR-ELISA

## Abstract

**Introduction:** Tuberculosis caused by *Mycobacterium tuberculosis* shows resistance to anti-tuberculosis drugs like Rifampicin and Isoniazid generating a public health problem. Using molecular techniques, such as PCR-ELISA, is an alternative that will detect resistance to RIF easily and economically.

**Objective:** Optimize a PCR-ELISA that can detect resistance to RIF in sputum samples.

**Method:** A PCR-ELISA was standardized to amplify a 255bp of rpoB gene that encodes resistance to RIF. Different parameters were optimized: hybridization temperatures, and probe concentration, among others. The technique’s analytical sensitivity, specificity, and concordance against GenotypeMTBDRplus v2 were evaluated using 20 samples in a pilot assay.

**Results:** A PCR-ELISA was optimized for the detection of resistance mutations; the analytical sensitivity of the PCR-ELISA was 4ng of PCR product for the H445D and H445Y probes while for the S450L probe the sensitivity was 0.4 ng of PCR product respectively. The technique has a specificity of 100%. Two mutations, S450W and L452P, were not detected by our system. In the pilot assay, a “good agreement” (k=0.737) was obtained between both techniques.

**Conclusions:** A PCR-ELISA was standardized for the detection of the 3 more frequent mutations in Peru associated with resistance to RIF simultaneously, the most frequent being S450L.

## 1 INTRODUCTION

Tuberculosis (TB) is caused by the bacillus *Mycobacterium tuberculosis* (MTb), and is transmitted when sick people expulse bacteria into the air (e.g., by coughing) (1) The disease typically affects the lungs (pulmonary TB) but can affect other organs of the body.

TB is a communicable, curable, and preventable disease; approximately 85% of people who develop the disease can be successfully treated (2). However, social determinants such as poverty, malnutrition, HIV infection, smoking, and diabetes aggravate this disease (2).

About 10 million people fall ill with TB and 1.4 million people die annually from the disease worldwide (3). Before the coronavirus pandemic (COVID-19), TB was the leading cause of death from a single infectious agent, ahead of HIV/AIDS (2). For the first time in more than a decade, TB deaths have increased due to reduced access to TB diagnosis and treatment and have reversed the progress made toward reaching the End TB Strategy targets set for 2030 and 2035 (2,3).

Drug-resistant TB remains a public health problem of great concern due to simultaneous resistance to isoniazid (INH) and rifampicin (RIF) the two most effective first-line drugs for TB treatment, which is called Multidrug-resistant TB (MDR-TB). Extensive drug-resistant (XDR-TB) tuberculosis which defined as a case of MDR-TB that is resistant to either fluoroquinolone and one of the second-line injectable drugs (capreomycin, kanamycin, or amikacin) (4) also contribute to worsening the problem. RIF-resistant TB (RR-TB), MDR-TB, and XDR-TB require longer treatment with second-line drugs causing more side effects when patients are treated (2).

Peru leads MDR-TB incidence cases in the Americas region unlike Brazil, which has a higher incidence of TB and HIV-TB cases (3).

In 2021, Peru reported 82 cases of tuberculosis with extensive resistance (XDR-TB) and 1256 cases of MDR-TB. 699 (55.7%) of these cases were reported in Lima and Callao, indicating a decrease compared to 2020, which reported 1205 MDR-TB cases, 664 (55%) of which were from Lima and Callao with 51 XDR-TB cases (5).

Rifampicin is a drug used for TB treatment. Resistance to this drug is caused by mutations in the 81 bp region (Hot Spot) of the rpoB gene, the 95% of the mutations that confer resistance to this drug are in this region and occur between codons 426 to 452 and generally consist of non-synonymous point mutations (6). In RIF-resistant isolates, mutations in codons 450, 445, and 430 of rpoB are the most frequent around the world, generating a high level of resistance however, the frequencies of single nucleotide polymorphisms (SNPs) in these three codons were variable in different geographic regions (7).

In Peru, the D435V, H445Y, H445D, and S450L changes are of high frequency generating resistance to RIF (8). Of these changes, S450L is the most frequent both in Peru and worldwide (9–11).

There are numerous microbiological methods to determine MTb resistance. Bactec Mycobacterium Growth Indicator Tube (MGIT) (12), Microscopic Observation Drug Susceptibility (MODS) (13), and Griess test (14) are good examples. The main problem with these techniques is that they take a long time to give the results. Nevertheless, currently exist molecular methods such as Xpert MTB/RIF (15) or GenotypeMTBDRplus v2 which are more sensitive, specific, and faster (16,17).

The WHO recommends the use of the Xpert MTB/RIF method because of its simplicity for rapid diagnosis of Rifampicin resistance, but this is an expensive method and is not capable of assessing INH resistance.

Another molecular technique called Genotype MTBDrplus v2 allows the detection of INH resistance genes (inhA and katG) and RIF genes (rpoB). Peru adopted the use of this technique due to its capability to detect both resistance genes and the high rate of monoresistance to RIF and INH (17,18).

The drawback of Genotype MTBDRplus v2 is the requirement for better infrastructure, expensive equipment, and highly trained professionals, and therefore they are difficult to implement in primary care hospitals and health centers, except in research centers. In countries with a high prevalence of TB such as Peru, the cost of supplies plays a decisive role in the adoption of some molecular techniques to detect TB resistance.

The PCR-ELISA technique, which combines a conventional PCR and an enzyme-linked immunosorbent assay (ELISA), was described more than 20 years ago (19) and has been used for the detection of many pathogens. There are many publications describing PCR-ELISA for the detection of foodborne pathogens such as *Salmonella tiphy, Listeria monocytogenes*, and *Vibrio parahaemolyticus* (19). Other publications have shown that PCR-ELISA was successful in the detection of gene resistance in other microorganisms (20–22). However, there is little information on the use of PCR-ELISA in tuberculosis, of the few that exist one describes the technique for predicting susceptibility to RIF of the *M. tuberculosis* complex (23) and a recent publication used the PCR-ELISA technique for the detection of resistance to INH and RIF, using MTB strains (24).

Generally, countries with high rates of tuberculosis are low-income countries or emerging economies that create dependency on foreign technologies. To cut this dependency, it is necessary to search for and develop methodologies that become diagnostic test that has the sensitivity, specificity, and speed of molecular methods, whose cost is like microbiological methods and that can mainly be implemented in primary care hospitals or less complex health centers. In this sense, this work aimed to optimize and evaluate, in the laboratory, a PCR-ELISA for the detection of H445D, H445Y, and S450L mutations conferring RIF resistance in sputum samples from tuberculosis patients.

## 2 MATERIALS AND METHODS

### 2.1 Sample collection and DNA extraction

The following strains were used for this work: ATCC N° 27294 (H37Rv) which was used as a control sensitive to all drugs, ATCC N° 35838 used as a control for resistance to RIF with the S450L mutation, and 2 strains (RIF-8853 and RIF-4469) of *Mycobacterium tuberculosis* which were used as controls for the H445D and H445Y mutations, from the National Reference Laboratory of Mycobacteria (LRNM) of the Peruvian National Institute of Health (INS).

For the pilot assay (In-laboratory evaluation), sputum samples were used. These samples were submitted to the LRNM for resistance determination by GenotypeMTBDRplus v2 (Hain Lifescience, GmbH) as a routine activity in the laboratory. From them, twenty remanent sputum samples with knowing resistance and 15 sensitive samples were used to evaluate the PCR-ELISA.

The sputum samples were provided by the LRNM and had the following characteristics: a bacilloscopy (BK+) ranging from 1+ to 3+ and a storage time in refrigeration (2 - 8°C) between 6 and 30 days since they were collected. The sensitive sputum was used for the CUT-OFF PCR-ELISA set-up.

Cetyltrimethylammonium bromide and lysozyme (CTAB-lyz) were used for DNA extraction (25,26): 200 μL of sputum sample was mixed with 40 μL of lysozyme (20mg/ml) in a 1.5 mL tube and incubated overnight at 37°C. Then 70 μL of SDS (10%) and 20 μL of proteinase K (10mg/mL) were added, vortexed, and incubated at 65°C for 40 min. Next, 100 μL of 5M NaCl and 100 μL of 10% CTAB were added to the tube, the sample was vortexed until the solution became milky white and then incubated for 10 min at 65°C. Subsequently, 750 μL of a chloroform/isoamyl alcohol solution (24:1) was added to the homogenate and centrifuged at 14 000 RPM for 5 min.

The aqueous phase was transferred to another sterile 1.5 ml tube taking care not to drag the interphase, 500 μL of ice-cold 100% Isopropanol (−20°C) was added, mixed by inversion 20 times, and then centrifuged for 15 min at 14000 RPM and the supernatant was removed. To the precipitate, 1 mL of ice-cold 70% ethanol was added, mixed by inversion 20 times, and centrifuged for 5 minutes at 14000 RPM, the supernatant was discarded, and the tubes were dried at room temperature. Finally, the DNA was resuspended in 200 μL of TE buffer and stored at −20°C until use.

All sputum samples were sent to the National Reference Laboratory of Mycobacteria (LRNM) to detect RIF and INH resistance as part of compliance with the Peruvian Technical Health Standard for Tuberculosis Control (NTS-041-MINSA/DGSP-V.01) (27). This document points out that all health centers must send sputum samples to LRNM to determine drug-resistant profiles.

The present study is part of two research projects reviewed and approved by the Institutional Research Ethics Committee of the Peruvian National Institute of Health (OI-057-14 and OT-003-20).

### 2.2 Primer and probe design

The primer sequences used in this work were obtained from Garcia et. al (23), the reverse primer (RP8T) had a digoxigenin (DIG) tag added at the 3’ end. The resistant probes for following mutation detection: H445Y, H445D, and S450L were designed from the 81bp sequence of the rpoB gene using the PrimerQuest™ program (IDT, USA) and were labeled with biotin at the 5’ end. Both were between 22 to 24 nucleotides in size. The primers and the labeled probes were synthesized by Eurofins™ (Table 1).

**Table 1.**
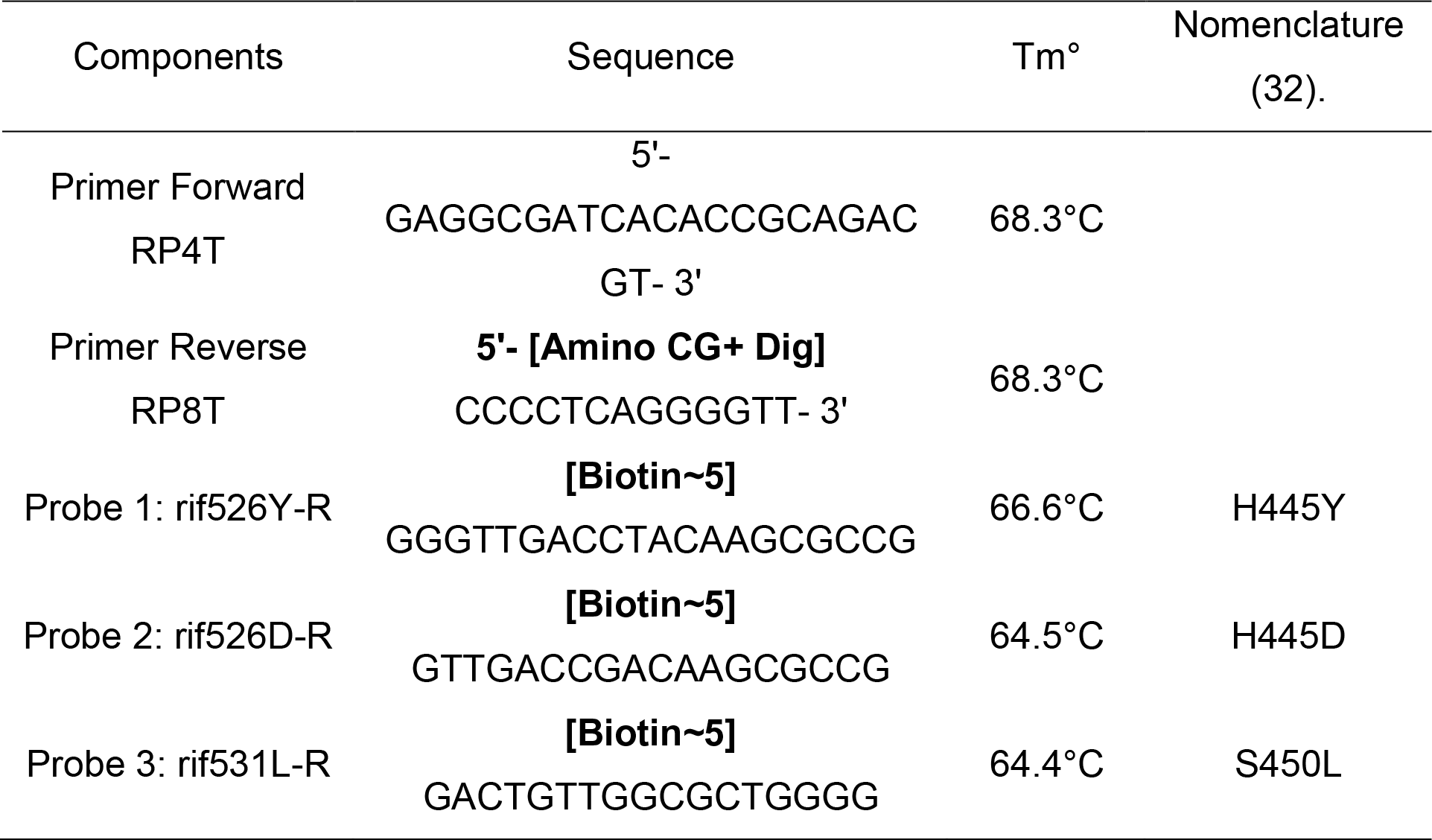
Sequences of primers and probes. The sequences of the RP4T and RP8T primers that amplify the 255 bp rpoB region and the sequences of the 3 resistant probes used in the pilot assay are shown.

### 2.3 Standardization of the PCR-ELISA

For the standardization of the PCR-ELISA, first, a PCR methodology was established to amplify a 255bp segment containing the 81 bp hot spot region of the rpoB gene. The different parameters were standardized such as the annealing temperature of the primers (60°C to 65°C) and magnesium chloride concentration (0 mM, 1 mM, 1.5 mM, 2 mM, and 3 mM).

For the ELISA, the first parameter to evaluate was hybridization temperatures between the PCR product and the probe at the following temperatures: 42°C, 45°C, 48°C, 50°C, 52°C, 55°C, and 58°C. Then, different degrees of stringency of the hybridization buffer (SSPE) were tested ranging from: low (3X), medium (1.5X), and high (0.5X). Different concentrations of streptavidin were evaluated: 1, 2, 4, 8, 10, 20, 40, 60 and 80 μg/mL, likewise different concentrations of probe were evaluated too: 0.5, 1, 5, 10 and 20 pmol/mL. The absorbance of the colored product of the ELISA was done at 450 nm.

Different concentrations of PCR product were also evaluated: 0.25, 0.5, 1, 1, 2, and 4 ng, finally, different dilutions of peroxidase-labeled Anti-DIG conjugate (AntiDIG-POD) were tested: 1/1000, 1/2000, 1/2500 and 1/3000.

To determine the analytical sensitivity of the PCR-ELISA, 2 dilutions of PCR product were performed: 1/10 and 1/100, starting from a concentration of 4ng for the 3 resistant probes. To determine the specificity of the PCR-ELISA, DNA from 7 Non-tuberculosis mycobacteria (NTM) determined by Genotype CM kit (Hain Lifescience, GmbH) were used: *Mycobacterium fortuitum, Mycobacterium malmoense, Mycobacterium avium, Mycobacterium chelonae, Mycobacterium kansasii, Mycobacterium intracellulare*, and *Mycobacterium abscesus*. In addition, human DNA was used as a negative control and 2 positive controls: H37Rv and ATCC strain No. 35838.

It is worth mentioning that for each parameter of the standardization, 4 controls were used, which were culture samples from resistant strains (RIF-8853, RIF-4469, and ATCC 35838) and a sensitive strain (H37Rv). Once the methodology was established, the PCR-ELISA was evaluated in a pilot assay.

### 2.4 Sequencing of a 255bp region of the rpoB gene

A 255bp region of the rpoB gene containing the 81 bp Hot Spot region associated with RIF resistance was sequenced from control samples ATCC N°35838, RIF-8358, and RIF-4469 to identify and verify the H445Y, H445D, and S450L mutations. From the purified PCR product, the sequencing reaction was performed using the BigDye™ Terminator v3.1 kit (Applied Biosystems), the sequencing reaction products were purified with the DyeEx 2.0 Kit (QIAGEN) and subsequently placed on the 3500 XL Genetic Analyzer (Applied Biosystems).

Samples that showed “inferred resistance” by GenotypeMTBDRplus v2 were also sequenced to identify mutations conferring resistance to RIF.

### 2.5 PCR-ELISA Pilot Assay (In-laboratory evaluation)

To establish the PCR-ELISA cut-off, DNA was extracted by the CTAB-lyz method from the 15 sensitive sputum samples and the H37Rv strain, then the PCR-ELISA was performed using all the parameters established in the optimization of the technique.

The CUT-OFF was determined using SPSSv25 software which allowed the calculation of the cut-off limit of the average of the absorbances of the 15 sensitive and H37Rv samples plus 3 times standard deviation (SD) for the H445Y and H445D probes and 2 times the SD for the S450L probe. A value above the CUT-OFF was considered resistant (presence of the mutation) and below was considered sensitive (absence of the mutation).

Once the cut-off values were obtained, the pilot assay was performed in blind (without knowing the resistance profile) using 20 sputum samples, whose resistance was previously determined by GenotypeMTBDRplus v2. DNA was extracted from these samples and PCR-ELISA was performed using the 3 probes simultaneously. The PCR-ELISA results were compared with the results obtained against GenotypeMTBDRplus v2 after revealing the blind.

### 2.6 Concordance analysis (Cohen’s Kappa Index)

The GenotypeMTBDRplus v2 method was chosen as a reference against PCR-ELISA because both techniques are molecular and allow the determination of mutations that confer resistance to RIF. The concordance between both techniques was determined using the following formula: K = Po − Pe/1 − Pe, where Po = is the proportion of observed concordance, Pe = is the proportion of concordance expected by chance, and 1 − Pe, represented the maximum possible agreement or concordance, not due to chance. The values of K can be found between zero and one, the closer to one the greater the concordance (28), the formula is implemented in the SPSSv25 software.

## 3 RESULTS

### 3.2 Standardization of PCR-ELISA

For annealing temperature of the primers and the magnesium ion concentration were established at 60°C and 1.5mM respectively, parameters at which a single amplification band of 255 bp was obtained (Table 2).

**Table 2.**
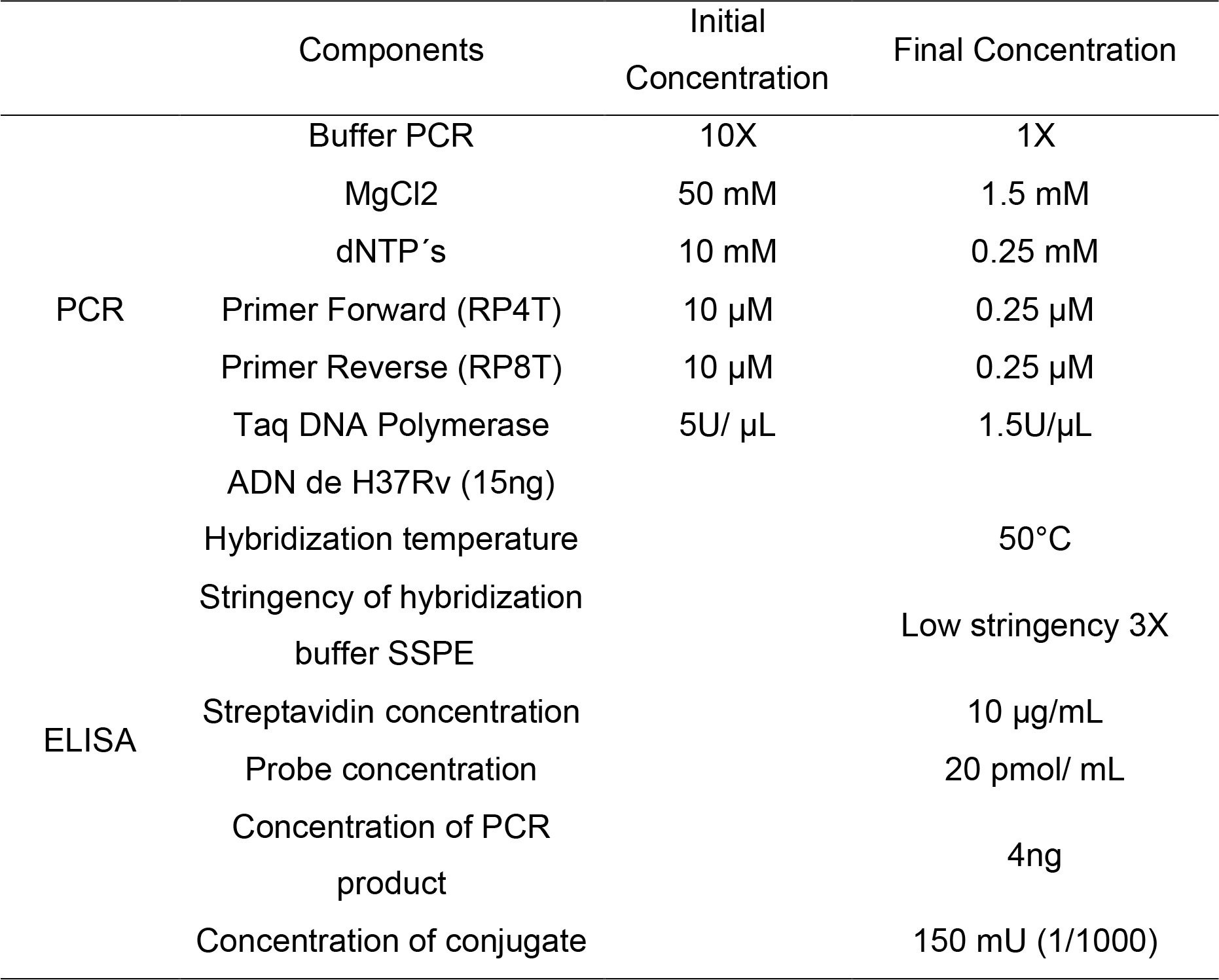
Optimized PCR-ELISA parameters. PCR-ELISA components and their different final concentrations after optimization of the technical parameters are shown

The PCR profile was established: Initial denaturation at 95°C for 10 min (1 cycle), denaturation at 95°C for 30 sec, annealing at 60°C for 45 sec and extension at 72°C for 30 sec (35 cycles). Final extension at 72°C for 5 min (1 cycle). PCR products were stored at −20°C until use.

Standardization of all PCR-ELISA conditions allowed to establish the following parameters: hybridization temperature at 50°C, SSPE hybridization buffer at low stringency (3X), streptavidin concentration at 10 μg/mL, probe concentration at 20 pmol/mL, PCR product concentration at 4ng and conjugate concentration at 150mU (1/1000) (Table 2).

### 3.3 Analytical Sensitivity of the PCR-ELISA

The analytical sensitivity of the PCR-ELISA was 4ng of PCR product for the H445D and H445Y probes while for the S450L probe the sensitivity was 0.4 ng of PCR product (Table S1).

### 3.4 Specificity of the PCR-ELISA

In the evaluation of the PCR-ELISA applied to *Mycobacterium fortuitum* and *Mycobacterium chelonae*, it was observed that there was no amplification of the 255 bp region; unlike the rest of the NTM (Figure 1) that did amplify the band of interest in addition to other non-specific bands.

**Figure 1.**
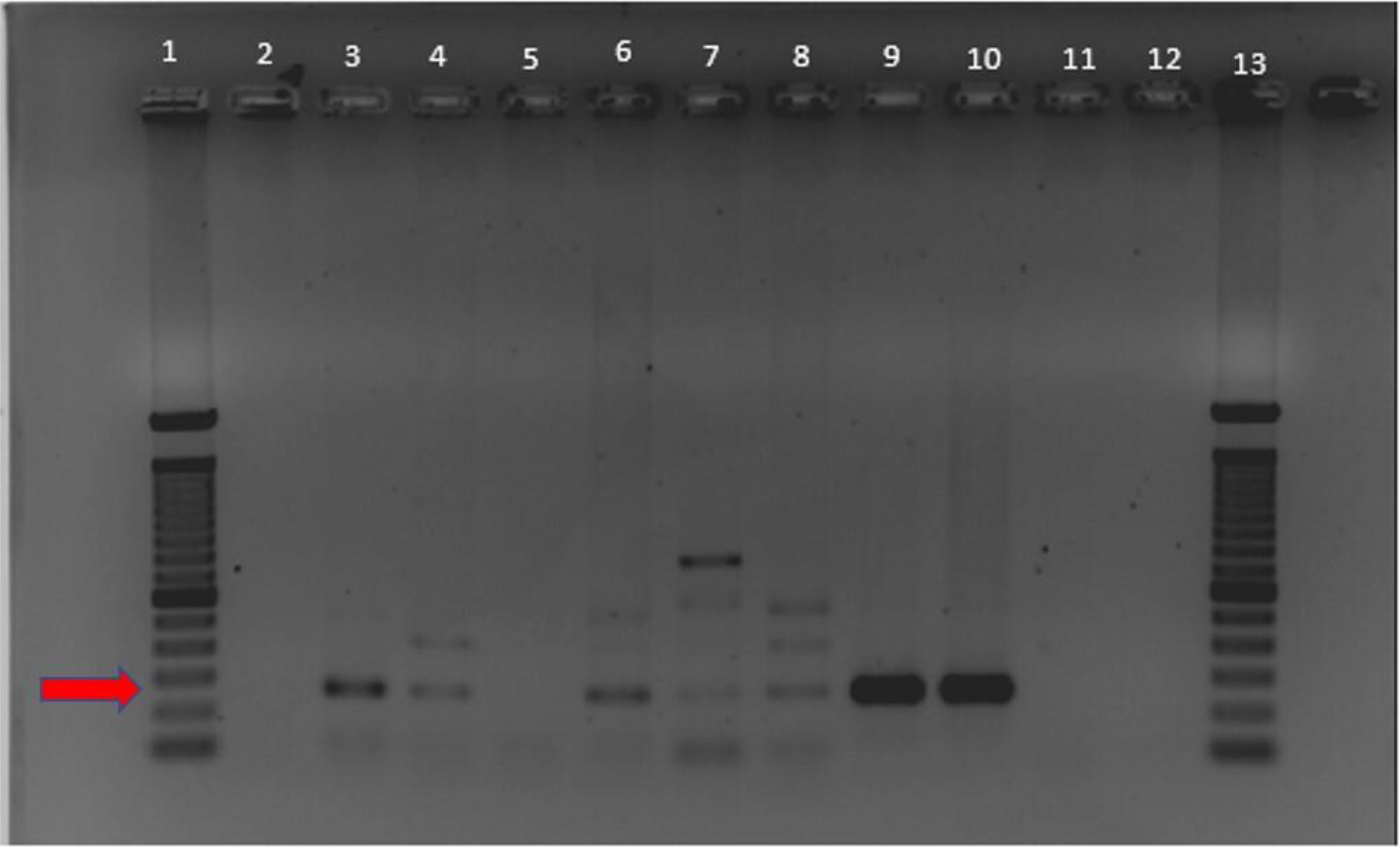
PCR-ELISA specificity. This 1.5% agarose gel shows the amplification product of 255bp of the rpoB region. Lane 2. Mycobacterium fortuitum. Lane 3. Mycobacterium malmoense. Lane 4. Mycobacterium avium. Lane 5. Mycobacterium chelonae. Lane 6. Mycobacterium kansasii. Lane 7. Mycobacterium intracellulare. Lane 8. Mycobacterium abscesus. Lane 9. H37Rv (positive control). Lane 10. ATCC 35838 (positive control). Lane 11. Human DNA (negative control). Lane 12. System control and Lane 1 to 13 (100 bp marker).

For the samples that did amplify, the PCR-ELISA gave negative results, determining that there was no hybridization in the NTMs nor the negative control (Table S2). The positive controls (C+) for the 3 probes hybridized correctly. Finally, the PCR-ELISA only detected *Mycobacterium tuberculosis*.

### 3.5 Sequencing of the 255bp rpoB region

The sequences obtained were analyzed using the online program: CLUSTAL W (https://www.genome.jp/tools-bin/clustalw) to align the forward and reverse sequences of each sample until a consensus sequence was obtained which was translated from nucleotides to amino acids using the ExPASy program (https://web.expasy.org/translate/). The sequences were visualized using the BioEdit program.

The analysis identified the presence of the following mutations: In sample RIF-8853 the nucleotide changes C → G was located at position 1335 which caused the substitution of amino acid Histidine by Aspartate at codon 445. For sample RIF-4469 the nucleotide changes C → T were observed at position 1335 and caused the amino acid variation Histidine by Tyrosine. For strain ATCC No. 35838 the nucleotide changes C → T were located at position 1335 and generated the amino acid substitution Serine for Leucine at codon 450.

### 3.6 PCR-ELISA Pilot Assay

The CUT-OFF of the system was established for the 3 resistant probes: H445D, H445Y, and S450L. The S450L probe had a cut-off of 0.641 and for the H445D and H445Y probes, the cut-off was 0.054 and 0.413 respectively (Table S3). Only for the H445D probe, it was decided to establish the CUT-OFF at 0.1 absorbances because the positive control had a high value (OD=0.426) which allowed establishing a better CUT-OFF and thus avoiding any false resistance in the samples analyzed.

When the blind analysis of the pilot assay was revealed, 3 samples did not amplify (M-4, M-5, and M-18) and therefore were not considered in the PCR-ELISA analysis (Table 3). Of the remaining 17 samples, 2 samples were not considered (M-11 and M-13), because these samples presented the absence of the Wild Type 8 (WT8) signal in the GenotypeMTBDRplus v2 (called “inferred resistance”) and which were not detected by the resistant probes of the PCR-ELISA system.

**Table 3.** PCR-ELISA results of 20 samples analyzed by the pilot assay.The results of smear microscopy, DNA concentration, the absorbance of the 3 probes, and resistance profile between PCR-ELISA vs GenotypeMTBDRplus v2 are shown.

| Samples | No. of crossings per BK | Concentration of DNA | Absorbances (O. D=450nm) |  |  | Resistance profile by Genotype | Comparison of techniques |  |
| --- | --- | --- | --- | --- | --- | --- | --- | --- |
|  |  |  | H445D | H445Y | S450L | rpoB | GENOTYPE | PCR-ELISA |
| M1 | 2+ | 2.07 ng/μL | 0.044 | 0.153 | 0.649 | MUT 3 | Resistant | Resistant |
| M2 | 2+ | 0.78 ng/μL | 0.086 | 0.238 | 0.866 | MUT 3 | Resistant | Resistant |
| M3 | 3+ | 1.60 ng/μL | 0.05 | 0.177 | 0.486 | WT | Sensible | Sensible |
| M4 | 2+ | 1.40 ng/μL | 0.005 | 0.012 | 0.059 | WT | Sensible | ND |
| M5 | 1+ | 1.10 ng/μL | 0.012 | 0.022 | 0.119 | MUT3 | Resistant | ND |
| M6 | 2+ | 3.26 ng/μL | 0.016 | 0.087 | 0.211 | WT | Sensible | Sensible |
| M7 | 3+ | 1.58 ng/μL | 0.075 | 0.233 | 0.844 | MUT3 | Resistant | Resistant |
| M8 | 3+ | 1.35 ng/μL | 0.065 | 0.22 | 0.696 | * | Resistant | Resistant |
| M9 | 2+ | 0.085 ng/μL | 0.032 | 0.101 | 0.488 | MUT3 | **Resistant** | **Sensible** |
| M10 | 2+ | 3.03 ng/μL | 0.058 | 0.172 | 0.795 | MUT 3 | Resistant | Resistant |
| M11 | 1+ | 3.40 ng/μL | 0.03 | 0.127 | 0.147 | * | **Resistant** | **Sensible** |
| M12 | 2+ | 4.04 ng/μL | 0.041 | 0.152 | 0.646 | MUT3 | Resistant | Resistant |
| M13 | 1+ | 1.28 ng/μL | 0.013 | 0.096 | 0.146 | * | **Resistant** | **Sensible** |
| M14 | 1+ | 0.053 ng/μL | 0.018 | 0.074 | 0.284 | MUT3 | **Resistant** | **Sensible** |
| M15 | 1+ | 2.02 ng/μL | 0.075 | 0.262 | 0.816 | MUT3 | Resistant | Resistant |
| M16 | 1+ | 0.34 ng/μL | 0.042 | 0.15 | 0.598 | WT | Sensible | Sensible |
| M17 | 2+ | 0.99 ng/ μL | 0.01 | 0.056 | 0.206 | WT | Sensible | Sensible |
| M18 | 1+ | 1.47 ng/ μL | 0.008 | 0.032 | 0.101 | WT | Sensible | ND |
| M19 | 2+ | 7.5 ng/ μL | 0.025 | 0.182 | 0.449 | WT | Sensible | Sensible |
| M20 | 1+ | 6.2 ng/ μL | 0.025 | 0.187 | 0.421 | WT | Sensible | Sensible |
| **Controls (c+)** |  |  | 0.321 | 0.651 | 1.118 |  |  |  |

ND. Not determined WT. Wild Type (\*) absence of WT8 MUT3: S450L

Upon sequencing these samples both were found to have mutations with the following variations: Serine for Tryptophan (S450W) and Leucine for Proline (L452P) at codons 450 and 452 respectively (Table S4), and for that reason, these samples were not considered in the concordance analysis.

Finally, 15 samples were evaluated between both techniques: 7 were resistant and the remaining sensitive and were concordant with GenotypeMTBDRplus v2. Two samples (M-9 and M-14) were discordant (Table 3).

### 3.7 Concordance analysis (Cohen’s Kappa index)

The PCR-ELISA system presented a sensitivity of 86.7% which corresponded to a successful concordance of 13 out of 15 samples. The value of Cohen’s Kappa index was 0.737, which denotes a “good agreement” between GenotypeMDRTBplus v2 and PCR-ELISA (Table 4).

**Table 4.**
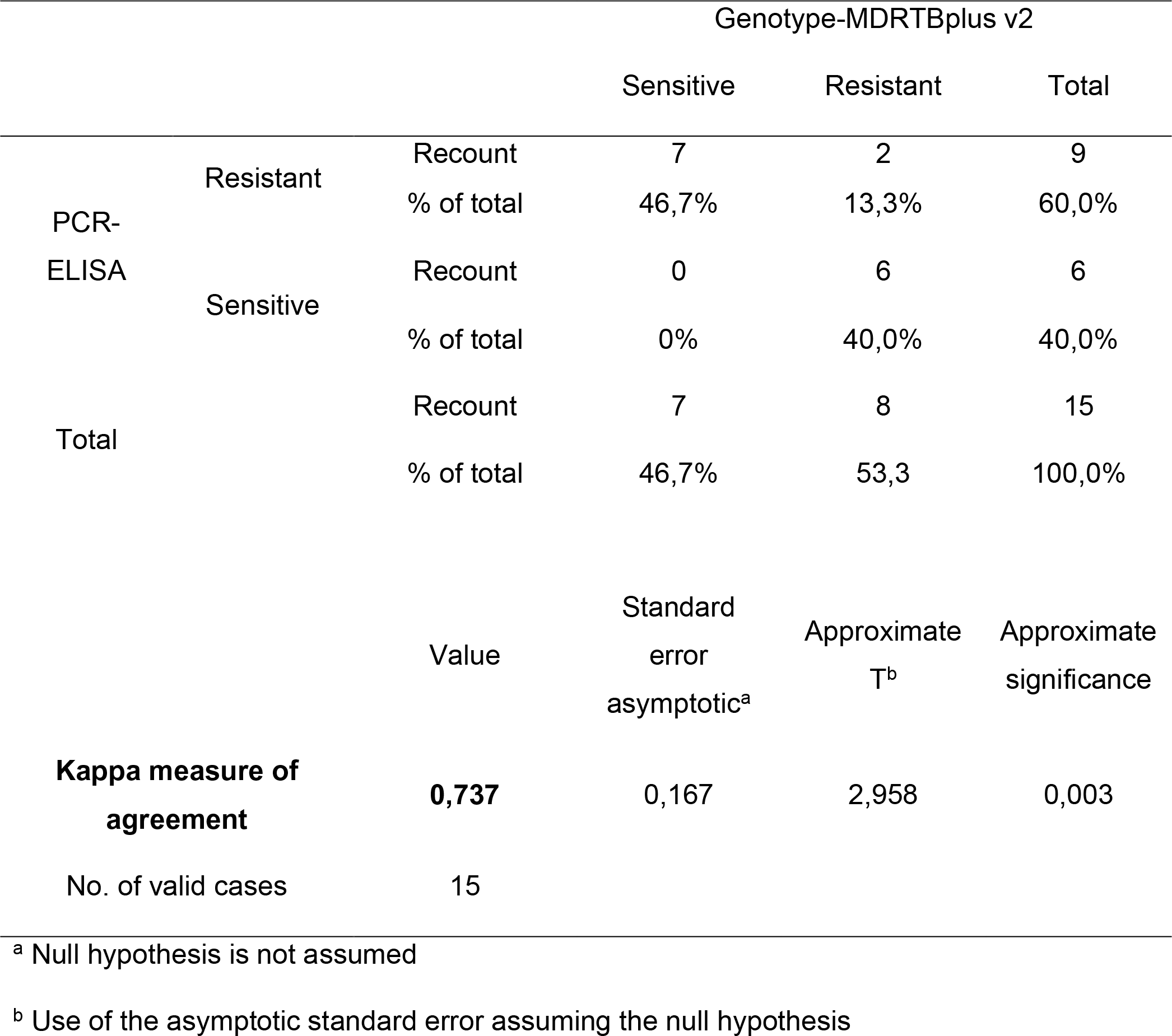
Cohen’s Kappa index between GenotypeMDRTBplus v2 and PCR-ELISA. The calculation of Cohen’s Kappa value was 0.737 which indicates a measure of “good agreement” between both techniques (N=15 samples).

## 4 DISCUSSION

This work presents the standardization and pilot evaluation of the PCR-ELISA technique for the detection of RIF resistance using biotinylated probes for which several parameters were optimized including DNA extraction.

In the present work, sputum samples and strains from which DNA was extracted by the CTAB-lyz method were used; this method proved to be adequate for direct extraction of DNA from sputum compared to other extraction protocols (29–31).

It was determined that the longer storage time of the samples and the lower number of crosses in the bacilloscopy may affect the concentration of DNA extracted from sputum (Table 3). A study by Ndhlovu et al. (29) where DNA was extracted from 40 samples with more than (2+) without further waiting time showed that they obtained good DNA concentration. Therefore, it would be important to extract DNA within 24 hours of sample collection, as prolonged refrigeration times possibly result in the lysis of cells and MTB bacilli.

For PCR standardization to amplify a 255 bp region of the rpoB gene, an annealing temperature gradient of the primers from 60°C to 65°C was performed. It was observed that different temperatures generate the same size amplification band. When observing the lane corresponding to human DNA, amplification of the band of interest was observed at all temperatures except at 60°C, so this temperature was chosen for PCR. This result was similar to that reported by Garcia et al. (23) who used the same primers, but whose PCR annealing temperature was at 65°C. The concentration of 1.5 mM MgCl2 was the most suitable as reported by Garcia et al. (23).

Sequencing results confirmed the point mutations in the 3 controls: ATCC 35838, RIF-8853, and RIF-4469 strains. The PCR-ELISA parameters were standardized (Table 2), in this technique we used the 3 resistant probes to detect RIF mutations that confer a high level of resistance and are also of high frequency in the Peruvian population (8,17,32). These mutations are in codons 450 (Serine for Leucine) and 445 (Histidine for Tyrosine or Histidine for Aspartate). These codons have also been shown to confer cross-resistance with rifabutin (10,33). It should be noted that rifabutin is one of the drugs in the rifamycin family recommended for the treatment of tuberculosis (TB) in HIV-infected persons during combination antiretroviral therapy containing HIV protease inhibitors (PIs), which reduces adverse effects (34).

The analytical sensitivity of PCR-ELISA for the H445D and H445Y mutations was 4 ng of PCR product (Table S1), while for the S450L mutation a sensitivity of 0.4ng was detected because the latter tends to have high absorbance values, this behavior coincides with that reported by Zhou et al. (24) and Garcia et al. (23) who evidenced different efficiencies between their probes. Probably the standardized parameters could have favored optimal conditions for a better hybridization of the S450L probe as opposed to the others, and therefore gave a better absorbance in the PCR-ELISA. Garcia et al. (23) showed that their PCR-ELISA system had a sensitivity of 0.1 pg in at least one of their five sensitive probes which in comparison with the PCR-ELISA system developed in this research was more sensitive, probably because they used sputum samples with 3 crosses while that our work evaluated samples from 1+ to 3+.

For the evaluation of the specificity of the PCR-ELISA, 7 NTM were used and of these 5 amplified and the other two *M. fortuitum* and *M. chelonae* did not (Figure 1). The samples that did not amplify were probably because the primers used did not recognize the region of interest. However, for the NTMs that did amplify when the ELISA was performed, they did not hybridize, therefore, the PCR-ELISA has 100% specificity (Table S2) confirming the specificity of the probes that recognize only *M. tuberculosis*. These results are similar to those obtained by Garcia et al (23).

During the establishment of the cut-off point, an adjustment had to be made to the CUT OFF value for the H445D probe, due to the reduced OD value (0.054), which did not allow a correct differentiation between resistant and sensitive samples. Therefore, due to the high OD value of its positive control, it was decided that the CUT OFF value should be 0.1, thus providing a better margin to establish the appropriate cut-off point and to standardize the discrimination criteria (S3).

This behavior of the probes confirms the fact that not all of them work with the same efficiency because the probes were designed and tested to identify those that give the best discrimination between the different alleles of the wild-type rpoB gene (23) and mutant or resistant alleles (24) respectively.

The PCR-ELISA results of the 20 samples showed that 3 of them (M-4, M-5, and M-18) had a BK between (1 to 2 +) and with a storage time in refrigeration between 6 to 20 days (Table 3), despite having a higher concentration of DNA there was no amplification of PCR product, probably because prolonged refrigeration times cause lysis in cells and MTB bacilli causing the DNA concentration detected during quantification to be more human DNA than bacillus DNA in these samples. Therefore, the PCR-ELISA analysis was performed on 17 samples (Table 3). Another finding of the pilot evaluation was that 2 samples (M-11 and M-13) were indirectly detected as resistant to RIF due to the absence of the Wild Type band (WT8) in the GenotypeMTBDRplus v2 between codons 447 to 452 (35), this situation is known as “inferred resistance” (8) and is because the resistance mutations are in another position and are not detected by the probes of the GenotypeMTBDRplus v2 reference method or by the probes used by the PCR-ELISA. These samples were sequenced to verify the presence of mutations outside the regions detected by both methods. In sample 11 the nucleotide change C → G was found at position 1350, which generated the substitution Serine for Tryptophan (S450W, Table S4.A), and for sample 13, the nucleotide change T → C was found at position 1356 which generated the variation of the amino acid Leucine for Proline (L452P, Table S4.B).

On the other hand, the samples (M-9 and M-14) were discordant in the detection of resistance. these samples had many days of refrigeration (approx. 24 days) and a bacilloscopy lower than 3+, this situation caused a low concentration of extracted DNA (<1ng/μL) which generated a faint amplification, poor hybridization, and therefore a false sensitive result in the PCR-ELISA (Table 3).

The concordance (kappa index “k”) between GenotypeMTBDRplus v2 and the PCR-ELISA technique was calculated. A value k = 0.737 was found (Table 4), which corresponded to a “good agreement” between both techniques.

In the total PCR-ELISA analysis based on 17 samples, when removing the 4 discordant samples: M-9 and M-14 due to low DNA concentration and M-11 and M-13 because the mutations were found outside the detection zone of the probes used, a value (k = 0.553) was obtained, which determined an “acceptable concordance” between both techniques.

Due to these findings, it was determined that the detection capacity of the PCR-ELISA will be affected when the samples have a BK lower than 3+ because there will be a lower number of bacilli, and the longer the number of days stored in refrigeration, the greater the possibility of lysis of the MTB cells. It was also observed that the sensitivity of the technique is affected when there are samples with mutations outside the detection zone of the probes used in this work. This research confirms what Garcia et al. (23) and Zhou et al. pointed out that the most predominant substitution associated with resistance to RIF was S450L. In addition, it was determined that this change was present in 7/13 samples (53.8 %). Finally, no samples with H445D and H445Y mutations were detected in the PCR-ELISA or the GenotypeMTBDRplus v2.

We report the use of PCR-ELISA as an alternative technique for the detection of rifampicin resistance directly from sputum samples, saving time in the determination of resistance. However, the technique has some limitations: detection of mutations for which no probes have been designed, e.g., D435V, cases of inferred resistance (absence of WT bands in GenotypeMTBDRplus v2), and samples whose mutation is in the same codon but encoding different amino acids (e.g., S450W and L452P).

Finally, it is necessary to validate the PCR-ELISA using a larger number of samples to determine the performance of the test with greater accuracy so that shortly, the test can be performed by health laboratories at the first level of care. In addition, the evaluation should be complemented by using probes for the detection of INH resistance and thus have a system that detects the 2 main drugs for the treatment of tuberculosis.

## 5 CONCLUSIONS

This work proposes the use of PCR-ELISA as an easy and economical alternative technique for the determination of RIF resistance directly from sputum, since, unlike other molecular tests, it does not require the use of expensive equipment such as those used in Genotype MTBDRplus v2, GeneXpert MTB/RIF, Real-time PCR or DNA sequencing. Instead, it uses basic, inexpensive equipment, and commonly available reagents, which are already present in most diagnostic laboratories. For these reasons, PCR-ELISA could be an alternative technique for inexpensive detection of RIF resistance and is easy to implement in any healthcare facility.

## ACKNOWLEDGMENTS

The authors are especially grateful to the National Reference Laboratory of Mycobacteria (LRNM) who supported us with the sputum samples and ATCC strains of *M. tuberculosis*. The authors also thank David García for his technical support in carrying out this study.

